# Modulation of kinesin’s load-bearing capacity by force geometry and the microtubule track

**DOI:** 10.1101/587089

**Authors:** Serapion Pyrpassopoulos, Henry Shuman, E. Michael Ostap

## Abstract

Kinesin motors and their associated microtubule tracks are essential for long-distance transport of cellular cargos. Intracellular activity and proper recruitment of kinesins is regulated by biochemical signaling, cargo adaptors, microtubule associated proteins and mechanical forces. In this study, we found that the effect of opposing forces on the kinesin-microtubule attachment duration depends strongly on experimental assay geometry. Using optical tweezers and the conventional single-bead assay we show that detachment of kinesin from the microtubule is likely accelerated by forces vertical to the long-axis of the microtubule due to contact of the single bead with the underlying surface. We used the three-bead assay to minimize the vertical force component and found that when the opposing forces are mainly parallel to the microtubule the median attachment duration between kinesin and microtubules can be up to 10-fold longer than observed using the single-bead assay. Using the three-bead assay, we also found that not all microtubule protofilaments are equivalent interacting substrates for kinesin and that the median attachment duration (median-Δt) of kinesin varies by more than 10-fold, depending on the relative angular position of the forces along the circumference of the microtubule. Thus, depending on the geometry of forces across the microtubule, kinesin can switch from a fast detaching motor (median-Δt < 0.2 s) to a persistent motor that sustains attachment (median-Δt > 3 s) at high forces (5 pN). Our data show that the load-bearing capacity of the kinesin motor is highly variable and can be dramatically affected by off-axis forces and forces across the microtubule lattice which has implications for a range of cellular activities including cell division and organelle transport.

**Significance Statement:** Kinesins are cytoskeletal motors responsible for the transport of cargoes along microtubules. It is well known that opposing forces decrease kinesin’s speed and run length. In this study, we found that when the pair of opposing forces applied on the kinesin-microtubule complex are parallel to the microtubule, the ability of kinesin to remain attached to the microtubule can vary by more than an order of magnitude depending on the relative azimuthal position of the pair of forces along the periphery of the microtubule. These results reveal a previously unknown versatility of kinesin’s load bearing capacity and as such have implications for the potential physiological roles of kinesin in a wide range of cell activities, including organelle transport and cell division.

---

Microtubules are cytoskeletal filaments essential for long-distance transport of intracellular cargos towards their plus and minus ends via kinesin and dynein motors, respectively (1). The stepping behavior of kinesin has been studied in detail at the single molecule level, both in the presence and in the absence of external loads on the motor (2–8). The effects of external forces parallel to microtubule on kinesin have largely been studied using the single-bead optical tweezers assay (2). Forces lateral to its travelling path have also been shown to affect kinesin’s processivity (9). Recent theoretical work suggests that an additional vertical force component due to contact of the single bead with the underlying surface has not been accounted for, and this force component must be considered when interpreting kinesin behavior acquired using this assay (10).

Vertical forces on kinesin are negligible when using a dual optical tweezers configuration and the three bead assay (11), but this assay has not been commonly used to study kinesin processivity. Additionally, the three bead assay can apply forces across the microtubule lattice with varied geometries (see below), which may be relevant, as it has been suggested that shear deformations applied to microtubules might affect kinesin activity (12). Indeed, recent experimental studies have shown longitudinal expansion of microtubules by 1.3 % to 1.6 % in the absence of external forces upon binding of saturating kinesin concentrations (13, 14).

To experimentally explore the effect of the direction of forces applied to the kinesin-microtubule complex, we used single-bead (Fig. 1A) and three-bead assays (Fig. 1C) to explore the role of vertical forces and forces across the microtubule lattice on kinesin activity. We found that vertical forces in the three bead assay accelerate kinesin detachment from microtubules. Additionally, we found that the median attachment duration of kinesin varies by more than 10-fold, depending on the relative angular position of the forces along the circumference of the microtubule.

**Fig. 1.**
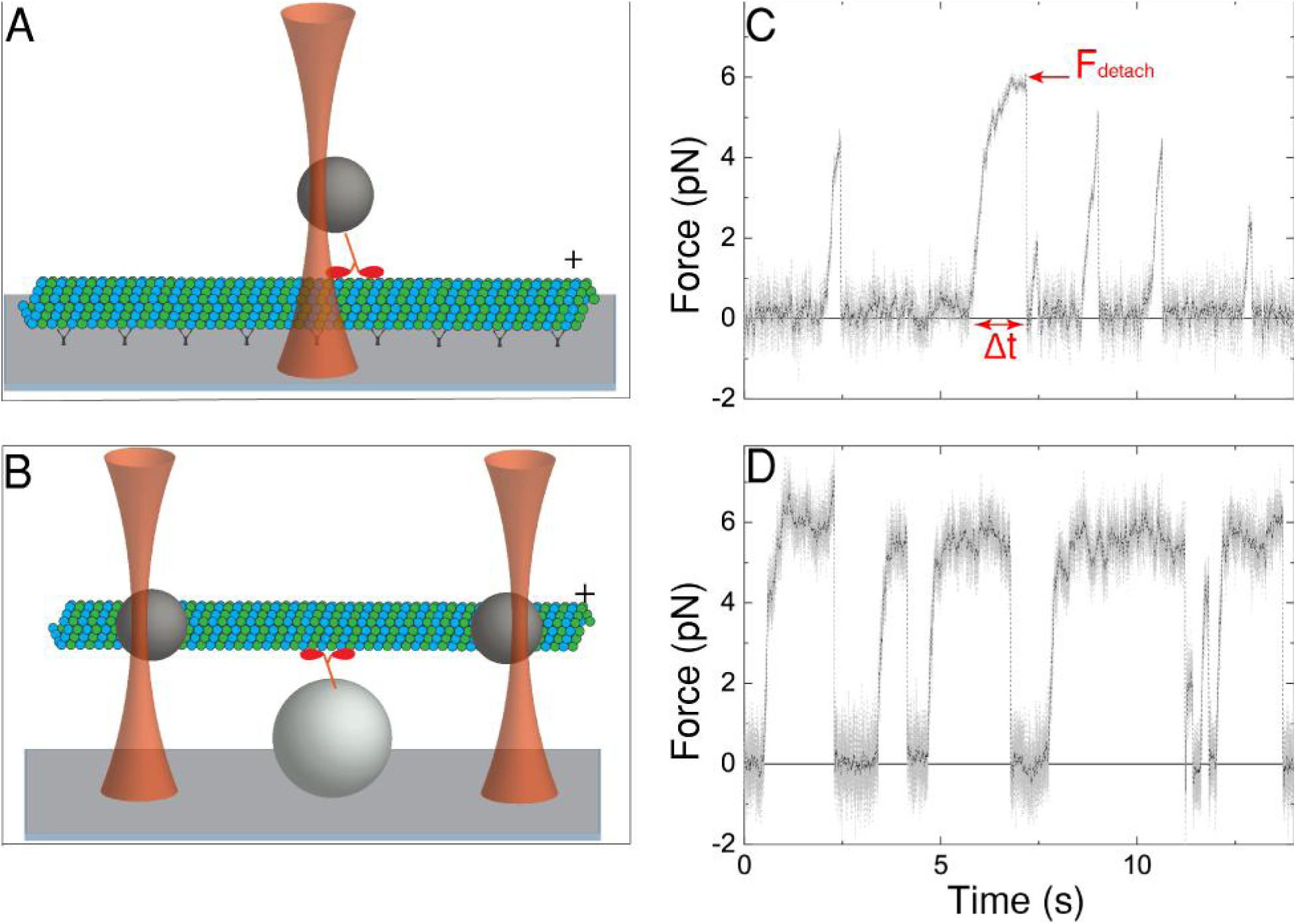
Single and three-bead assays for kinesin-microtubule interaction. Cartoon representations (not drawn to scale) of (A) the single-bead assay and (B) the three-bead assay. (A) A microtubule (green and blue) is immobilized on a glass coverslip (gray surface) via antibody or streptavidin (black). A kinesin molecule (red) attached on a bead (gray sphere) is brought into contact via optical tweezers (brown). (B) A microtubule is attached to two laser-trapped beads via streptavidin-biotin linkage. The microtubule-assembly (microtubule dumbbell) is brought in contact with a single kinesin molecule, attached on surface-immobilized spherical pedestal. Representative raw (gray) and smoothed (black) force traces of a single kinesin molecule interacting with (C) a surface-immobilized microtubule in the single-bead assay and (D) a microtubule dumbbell in the three-bead assay. The force (*F*_detach_) at which the kinesin detaches (red arrow in (C)) from the microtubule and the duration Δ*t* of the corresponding force ramp (double red arrow in (C)) can be calculated directly from the data for every force ramp.

## Results

### Kinesin attachment durations in the single-bead assay depend on bead diameter

A two-headed kinesin construct (17) was site-specifically bound to beads of three different diameters (2.1 μm, 0.82 μm and 0.51 μm, *Materials and Methods*), which were held in a stationary optical trap as the motor stepped along coverslip-attached taxol-stabilized GDP-microtubules. Processing kinesin pulled the bead out of the center of the trap, increasing the resisting force on the kinesin until the motor detached from the microtubule (Fig. 1A), resulting in the bead moving rapidly back to the center of the trap before the motor reengaged to start another run. The corresponding force traces were similar to those published previously with the same kinesin construct, ATP concentration, and bead diameter (15). For each processive run, the force component *F*_x_ at detachment (*F*_detach_) and the duration of attachment (Δ*t*) were measured (Fig. 1C).

We examined the effect of vertical forces on *F*_detach_ and Δ*t* by varying the bead diameter. The vertical *F*_z_ component experienced by the bead (10) is expected to scale with the radius R as 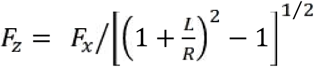 Eq. 1, where *F*_x_ is the force measured by the laser trap and *L* (∼35 nm) the length of the kinesin construct (Fig. 2A). The resulting measurements of *F*_detach_ and Δ*t* for the three different sizes of bead at 2 mM MgATP are plotted in Fig. 2B. The corresponding distributions of both Δ*t* and *F*_detach_ for the largest (2.1 μm) and smallest (0.51 μm) bead diameter are statistically significantly different (p < 0.001 Mann-Whitney test; n = 164 from 4 different beads and 166 runs from 4 different beads, respectively). As can be seen from the cumulative frequency distributions of Δ*t* and *F*_detach_ (Fig. 2B) kinesin detaches faster from the microtubule and reaches lower detachment forces when it is bound to the larger 2.1 μm rather than to the smaller 0.51 μm bead diameter (Table I). The frequency of stall events significantly decreases from 30% down to 5% and 0% with increasing bead diameter of 0.51 μm, 0.82 μm and 1.0 μm, respectively (Table I). The corresponding stall force values of 4.4 ± 0.81 and 4.9 ± 0.43 pN (SD: standard deviation) are close to previous measurements of 4.8 – 7.4 pN for kinesin 1 at saturating ATP (2, 3, 5, 15, 16). The average detachment force <*F*_detach_> of 2.9 pN ± 1.1 pN (SD), is also comparable with a previously measured value of 4.4 ± 1.4 pN (15) for the same kinesin construct and bead diameter (0.82 μm) and an estimated characteristic value of 3.2 pN (7).

**Fig. 2.**
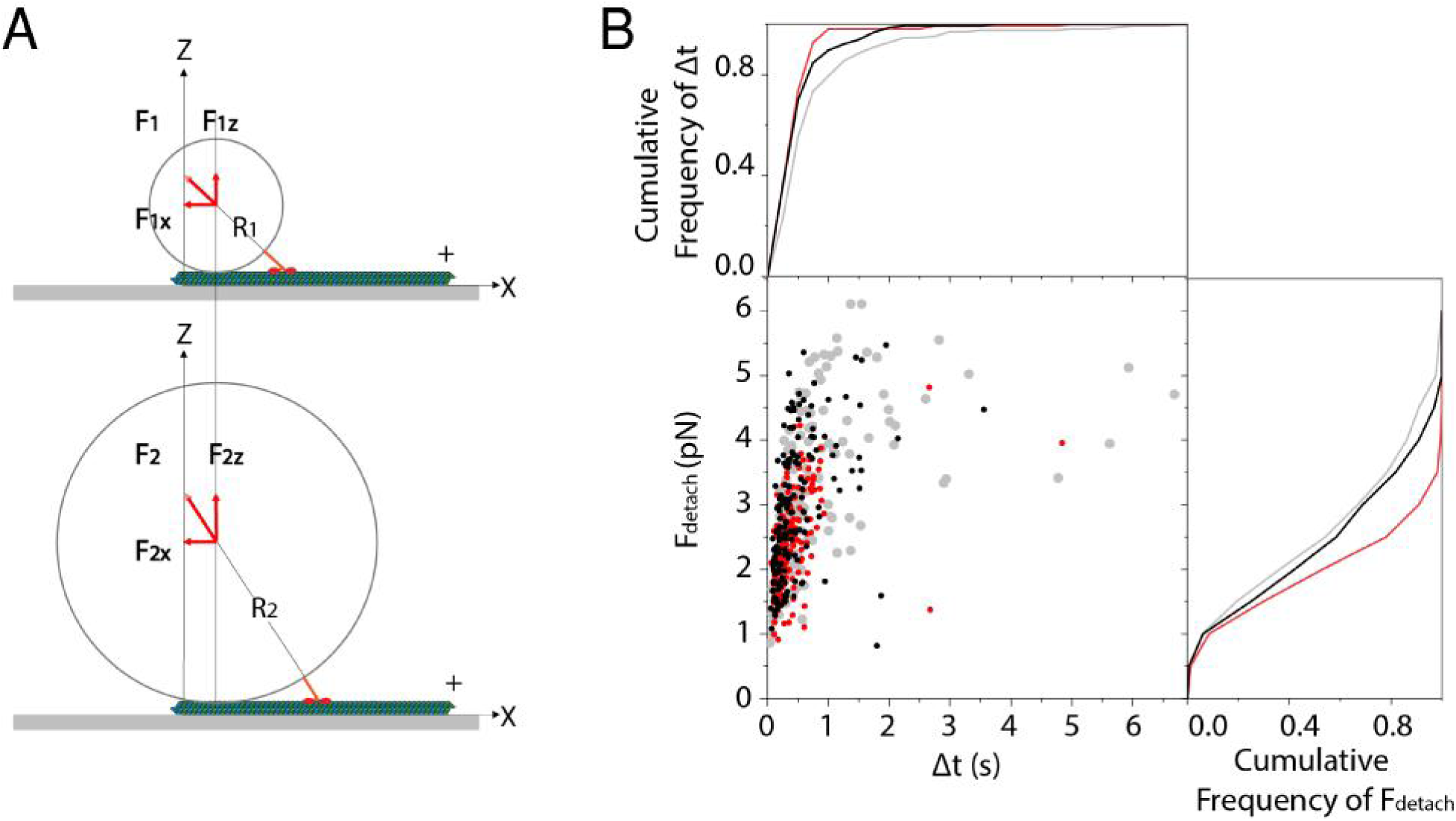
Kinesin detaches faster and at lower forces for larger size beads. (A) Cartoon representation of the single-bead assay with two different sizes of beads R_1_ < R_2_ and the same displacement along the microtubule relative to the center of the laser trap. The components (F_x_, F_z_) and the vectors of the total force F that opposes kinesin’s movement are indicated by the red arrows. Mechanical equilibrium dictates that F should be opposite to the force produced by kinesin and lay along the radius of the bead. For the same stiffness of the trap F_1x_ = F_2x_ and F_1z_ < F_2z_ (see Eq. 1). (B) Scatter plot of *F*_detach_ and Δ*t* for beads with three different diameters of 0.51 μm (gray), 0.82 μm (black) and 1.0 μm (red). The cumulative distributions of *F*_detach_ and Δ*t* are plotted as solid lines of corresponding colors along the right and top axis of the graph, respectively.

**Table I.**
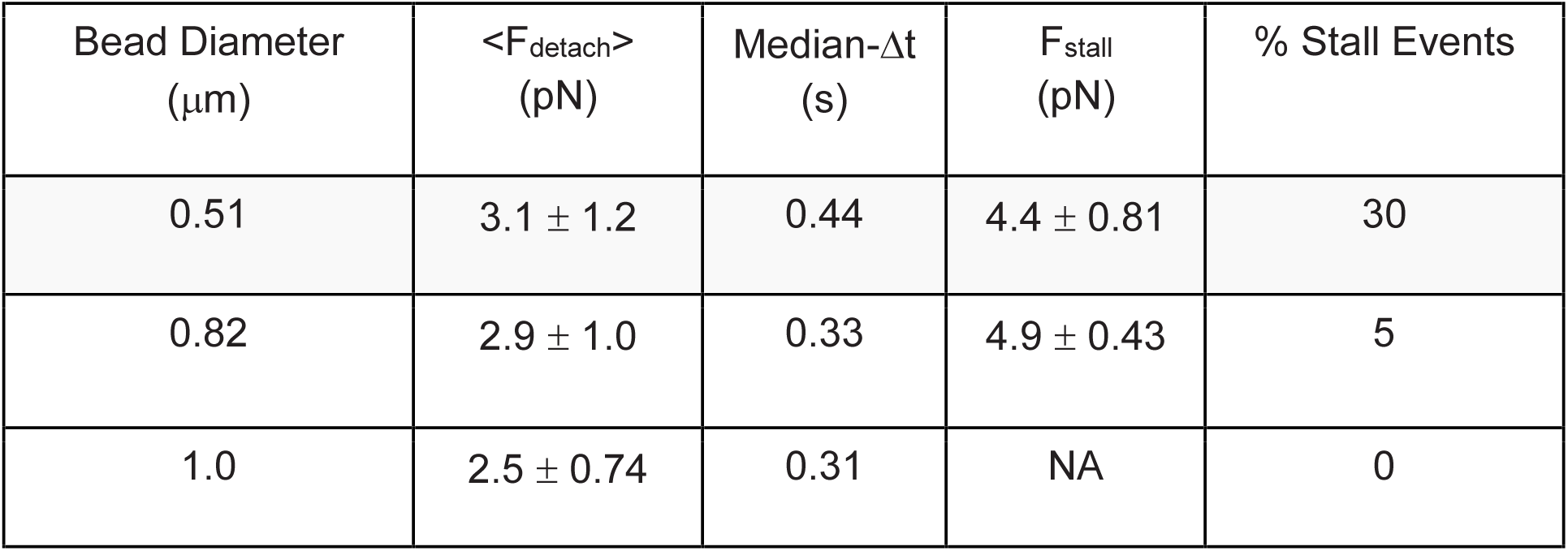
Single-bead assay measurements of forces and attachment durations.

| Bead Diameter<br>($\mu\text{m}$) | $\langle F_{\text{detach}} \rangle$<br>(pN) | Median-$\Delta t$<br>(s) | $F_{\text{stall}}$<br>(pN) | % Stall Events |
| --- | --- | --- | --- | --- |
| 0.51 | $3.1 \pm 1.2$ | 0.44 | $4.4 \pm 0.81$ | 30 |
| 0.82 | $2.9 \pm 1.0$ | 0.33 | $4.9 \pm 0.43$ | 5 |
| 1.0 | $2.5 \pm 0.74$ | 0.31 | NA | 0 |

We checked different substrates and/or microtubule surface-attachment strategies and found no effect on the mechanical activity of kinesin-1 stepping along microtubules attached (a) to a solid surface via tubulin antibody or via streptavidin or (b) to a biotinylated supported lipid bilayer via streptavidin (*Materials and Methods*). Representative measurements for different surfaces and attachments are shown in Fig. 3E and Fig 3I (Table S1).

**Fig. 3.**
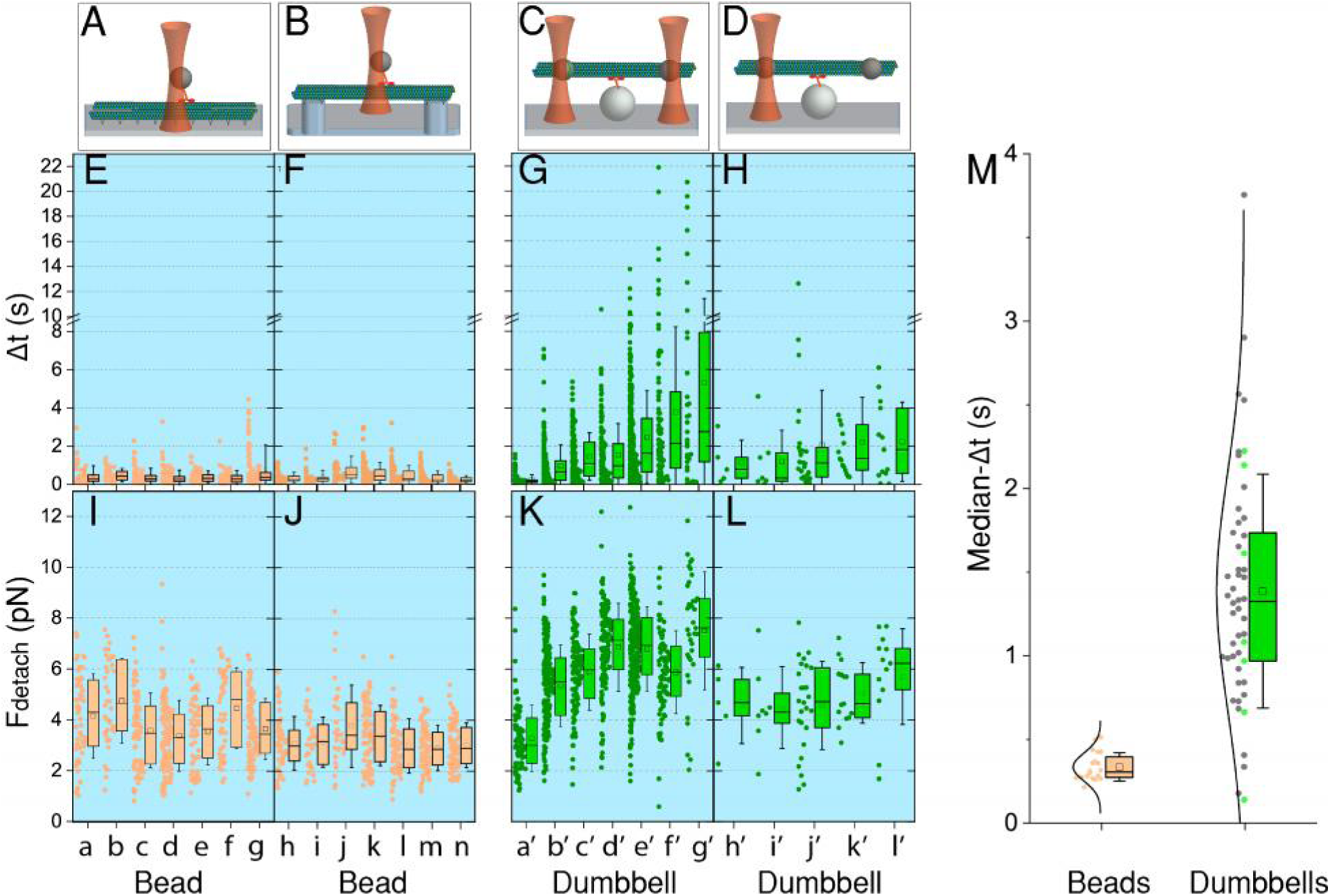
Kinesin’s attachment durations and detachment forces for single and three-bead assays. Cartoon descriptions for each assay are shown on plots (A), (B), (C) and (D). Representative distributions of Δ*t* and *F*_detach_ and their corresponding box-statistics for: (E, I) different pairs of beads (“a” to “g”) and surface immobilized microtubules (Table S1); (F, J) different pairs of beads (“h” to “n”) and microtubules suspended between rectangular ridges; (G, K) different microtubule dumbbells (“a’” to “g’”) and (H, I) microtubule dumbbells (“h’” to “I’”) laser trapped only by the bead at plus end and interacting with single kinesin. For each box the width corresponds to the inter-quartile range (IQR), the error bars to standard deviation (SD), the mid-line to the median value and the square inside each box to the average value. (M) The distribution and box statistics of the median-Δ*t* for single kinesin molecules interacting with microtubule dumbbells (n = 50) and surface-immobilized microtubules (n = 20) under resisting load. The green points correspond to the examples shown in plot (G). The black lines, which mainly serve as a guide to the eye, represent a normal distribution with the same mean and standard deviation as the corresponding data.

To test if the lack of attachments along the microtubule segment that interacts with kinesin may lead to different behavior, we fabricated parallel rectangular pedestal ridges (4 mm long, 2 µm wide, 1 µm tall and 10 µm apart; *Materials and Methods* and Video S1). Microtubules were immobilized across the top of the ridges (Fig. S1, Video S1) and the single bead assay along the part of microtubules suspended between rectangular ridges (Fig 3B) revealed similar values to those for the surface-immobilized microtubules (<*F*_detach_ > = 3.1 ± 1.9 pN (SD, n = 432 runs); (<median-Δ*t* >_w_ = 0.30 ± 0.11 s (SD_w_, n = 7)) (Fig. 3F & 3J).

### Kinesin exhibits a broad range of microtubule attachment durations in the three-bead assay

To apply mainly shear forces on the microtubule – kinesin complex and reduce the effect of the force component vertical to the microtubule axis, which is inherent to the single-bead assay (10), we used a dual laser trap and the three-bead assay (Fig. 1B). The three-bead assay has been used mostly for single molecule studies of the non-processive actomyosin complex (11, 17) and less frequently for microtubule interacting motors (18–20). Two streptavidin-coated polystyrene beads (dia. = 0.82 μm), trapped by two laser beams ∼10 um apart, were brought in contact with a taxol-stabilized biotinylated GDP-microtubule in solution until a stable “dumbbell” assembly was formed (Fig. 1B). Initial shear pretensile forces (*F*_pretensile_ = 2-9 pN) were applied to the dumbbells by moving the two laser beams further apart. Kinesin molecules were anchored on surface-immobilized spherical pedestals (dia. = 2.5 um) against which microtubule dumbbells were brought in contact. Single molecule kinesin motility resulted in the displacement of the dumbbells relative to the surface immobilized kinesin. Single kinesin molecules (Materials and Methods; Fig. S2) interacting with microtubule dumbbells in the presence of 2 mM MgATP reached forces ≥ 5 pN more frequently and remained attached at these high forces remarkably longer than observed in the single-bead assay (Fig 1D). Representative examples of the distribution and statistics of Δ*t* and *F*_detach_ between kinesin and different microtubule dumbbells are shown in Fig. 3G and Fig. 3K, correspondingly. The median duration of the runs produced by kinesin varied for different dumbbells by more than an order of magnitude (0.14 s and 3.8 s) and followed a unimodal symmetric distribution (Fig. 3M) with a weighted mean (Supporting Information) of <median-Δ*t* >_w_ = 1.3 ± 0.59 s (SD_w_: weighted standard deviation, n = 50). Single beads interacting with surface-immobilized microtubules didn’t demonstrate such a variability and the corresponding value was <median-Δ*t* >_w_ = 0.34 ± 0.083 s (SD_w_, n = 20) (Fig. 3M). To reduce the possibility that the large variability in attachment durations may be due to more than one kinesin molecules interacting with the microtubule dumbbells we examined the force plateaus for each dumbbell by measuring the corresponding stall forces (Supporting Information). The stall forces for each dumbbell were similar independent of median-Δ*t* (Fig. S3A). The average value of stall forces 6.8 ± 0.74 pN (SD_w_, n = 48 dumbbells) was well within the range of previously measured stall forces using the single-bead assay measurements 4.8 – 7.4 pN (2, 3, 5, 15, 16). The level of force plateaus during prolonged attachments was similar for all the kinesin runs within each dataset (Fig 1D) which lends additional confidence that the increased attachment durations are due to single kinesin molecules (10, 16, 21). Indeed for higher kinesin surface density approximately twice as high force plateaus could be observed (Fig. S3), in agreement with the prediction that when the vertical forces are small the stall force of two kinesins is twice that for a single kinesin (10). It is worth mentioning that similar recurring long attachments at similar high force plateaus ≥ 5pN between single molecule kinesin and microtubules at saturating ATP concentrations had been observed in the past for a microneedle assay that has similar geometry (3). The <*F*_detach_> for different dumbbells was between 2.8 pN and 8.0 pN depending on the frequency of kinesin runs that reach high forces before detaching from each dumbbell (Fig. 3K). During the initial motility phase (<*F*_detach_> ≤ 3 pN) the weighted mean of the average velocity of kinesin between the single-bead and three-bead assays differed by less than 10% (300 ± 65 nm/s vs. 280 ± 83 nm/s, respectively; Fig. S4). Similar differences between microtubule dumbbells and surface-immobilized microtubules, were observed when GMPCPP was used instead of GTP and taxol to produce stable microtubules (Fig. S5). This result suggests that the broad variability of median-Δ*t* observed in the three-bead assay (Fig. 3M) is not due to the variability in microtubule protofilament number (22, 23).

### The broad range of kinesin-microtubule attachment durations is not due to variations in pretensile forces or the tilting of the microtubule within the dumbbell

The distribution of Δ*t* between a single kinesin and a given dumbbell did not show dependence on the magnitude of the pretensile forces between 2 - 9 pN applied on the dumbbell (Fig. S6). Additionally, when no pretensile forces were applied and only the one bead attached to the plus-end of the microtubule was trapped (Fig. 3D), kinesin achieved the high forces (Fig. 3L) and long dwell times as observed for dually trapped dumbbells (Fig. 3H). Note that for most of the dumbbells <*F*_detach_> (∼ 6 pN) was greater than *F*_pretensile_ (∼4 pN). To test if the broad distribution of median-Δ*t* (Fig. 3M) is due to kinesins processing along microtubules not parallel to the coverslip (Fig. S7), we tested the sensitivity of the Δ*t*-distribution for a given dumbbell to changes in *z*-displacement. Increasing by 50 or 100 nm the separation along the z-axis between a given dumbbell and the kinesin changed only the probability of kinesin attachment, but not the Δ*t*-distribution (Fig. S8B, C, D, E). However, the z-force developed in the single bead assay is expected to be larger (∼ 12 pN, for R =0.41 μm, L = 35 nm and F_x_ = 5 pN; Eq. 1) than in the three-bead assay (∼0.4 pN; Fig. S7). This larger force has been proposed to accelerate the forced detachment kinetics of kinesin (10) and lead to a narrow distribution of median-Δ*t*, as we see in our experiments (Figure 3M).

### Kinesin-microtubule attachment durations depend on the relative azimuthal separation of opposing forces

The most variable experimental parameter among the dumbbells in the three-bead assay is the geometry of shear forces (Fig. S9). When microtubule dumbbells are formed, the trapped beads do not always bind at the same azimuthal position relative to each other or the surface-immobilized kinesin molecule. Therefore, the relative azimuthal positions of the pair of opposing forces between the interacting kinesin and the plus-end trapped bead will be variable in the three-bead assay (Video S2). If this angular variability is responsible for the broad distribution of attachment durations *Δ*t (Fig. 3M), then the values median-Δ*t*_1_ and median-Δ*t*_2_ of kinesin interacting at diametrically opposite positions along the cross section of the same microtubule dumbbell should be different.

To test this proposal, we collected two data sets for each dumbbell (n = 24 dumbbells). First the dumbbell was brought into contact with a pedestal on the bottom of the experimental chamber, and then to a pedestal on top of the same chamber (Fig. 4A). By testing this pair of interactions, we are probing diametrically opposite sides of the microtubule which is equivalent to rotating the microtubule around its axis, and thus changing the angle φ (Fig. 4B). We found the median-Δ*t*_1_ for pedestals on the bottom and median-Δ*t*_2_ for pedestals on the top to be anti-correlated with a Pearson coefficient of - 0.5 (p = 0.02). Note that the short and long interactions did not always occur on the same side of the chamber, but were rather random, arguing against a systematic artifact causing the anti-correlation (Fig. 4C). Indeed the distributions of median-Δ*t*_1_ and median-Δ*t*_2_ are similar to each other (Fig. 4D) and to the distribution in Fig. 3M.

**Fig. 4.**
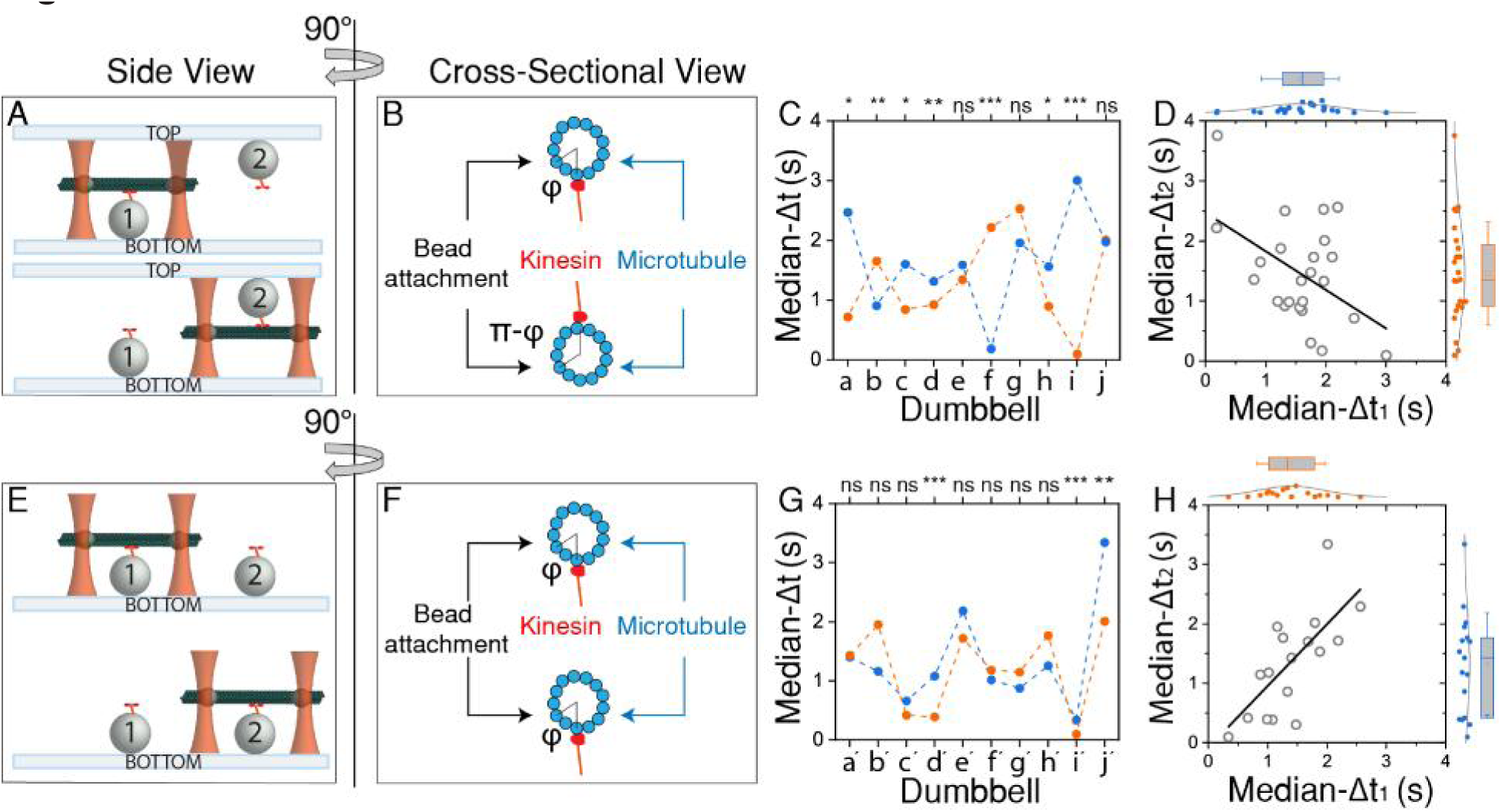
The interaction between kinesin and microtubule dumbbells depends on the relative azimuthal position φ of opposing forces. Cartoon representation of experiments in which the same dumbbell is brought against two different interacting pedestals “1” and “2” (A, B) in opposing surfaces (top and bottom) or (E, F) along the same surface. In (B) and (F) φ or π-φ indicate the relative azimuthal position between the attached bead and the interacting kinesin. Plots of median-Δ*t* for 10 different microtubule dumbbells against two different pedestals (Δ*t*_1_, Δ*t*_2_): (C) on the top and bottom surface and (G) along the same surface of the experimental chamber. Dumbbells “i” and “j” in plot (C) are the same dumbbells with “i′” and “j′”, respectively, in plot (G). The statistical significance of the difference between median-Δ*t*_1_ and median-Δ*t*_2_ for each dumbbell is shown on top of plots (C) and (G) (Mann-Whitney test; ns = not significant, * stands for p <0.05, ** for p<0.01, *** for p<0.001). (D) and (H) are scatter plots of median-Δ*t*_1_ and median-Δ*t*_2_ and solid lines represent linear fits. The distribution and statistics of median-Δ*t*_1_ and median-Δ*t*_2_ are plotted on the top and the right, respectively, of the plots (D) and (H).

To establish that the anti-correlation between median-Δ*t*_1_ and median-Δ*t*_2_ was due to a significant change of the relative azimuthal position, φ, we performed experiments this time testing the same dumbbell (n = 17) against two different pedestals on the same side of the experimental chamber (Fig. 4B and 4E). These experiments would reflect any variability due to differences in surface attachment orientations of kinesin and in its relative position with the interacting microtubule dumbbell. In contrast to the previous experiment, the Pearson correlation coefficient for median-Δ*t*_1_ and median-Δ*t*_2_ is positive 0.7 (p = 0.001) (Fig 4F). It is likely that the actual values of both correlation and anti-correlation coefficients are underestimated because we do not have a fine positional control at the protofilament level. The azimuthal position φ will not always be exactly the same for spherical pedestals across the same surface (Fig. 4F) and not always exactly π-φ for spherical pedestals in opposing surfaces (Fig 4B).

## Discussion

Our results show that the attachment duration and the magnitude of opposing forces that can be sustained by the microtubule-kinesin complex, or else kinesin’s load-bearing capacity, is highly variable and can be dramatically affected by the relative orientation between the force vector and the microtubule track. For different cellular processes in which the microtubule-kinesin complex can be subject to opposing forces such as during cargo transport and mitotic division the 3D orientation of the force vector can be significantly different between the two cases. A significant difference in the force vector due to the presence of a larger force component vertical to the microtubule in the single-bead assay relative to three-bead assay is most likely the reason for the observed differences in <median-Δ*t* > of the microtubule-kinesin complex between the two assays, supporting a recent theoretical study (10). Moreover, when opposing forces are oriented mainly along the microtubule axis (three-bead assay), kinesin’s load-bearing capacity depends on the relative angular position of the pair of opposing forces around the circumference of the microtubule. Depending on its location on the microtubule, kinesin can switch from a fast detaching motor to a persistent motor that can sustain its microtubule attachment at high forces (>5 pN) for extended lengths of time, with almost an order of magnitude difference. The mechanism for the observed variability of kinesin’s load-bearing capacity in the three-bead assay is likely related to the structural plasticity of the microtubule (24) and/or possible cooperative clustering of different post-translationally modified tubulin isoforms during microtubule polymerization, such that not all the protofilaments of a microtubule are equivalent interacting substrates for kinesin under opposing forces. Interestingly, recent structural studies have shown biochemically induced expansion and cross-sectional deformation of microtubules upon binding of molecular motors in the absence of external mechanical forces (14, 25). Nevertheless, our findings point to a more general mechanism by which the duration of the interaction between microtubules and binding partners can be a more complex function of the geometry of forces than previously appreciated.

## Materials and Methods

### Reagents

Guanosine Triphosphate (GTP) and paclitaxel (taxol) were purchased from Cytoskeleton Inc (Denver, CO, US). Guanosine-5’-[(α,β)-methyleno]triphosphate, sodium salt (GMPCPP), the non-hydrolysable analogue of GTP, was purchase from Jena Biosciences (Jena, Germany). piperazine-N, N′-bis (PIPES), Adenosine Triphosphate (ATP), glucose were purchased from Sigma. 4-(2-hydroxyethyl)-1-piperazineethanesulfonic acid) (HEPES) and Dithiothreitol (DTT) were purchased from GoldBio (St Louis, MO, US). Ethylene Glycol-bis(β-aminoethyl ether)-*N*,*N*,*N*′,*N*′-Tetraacetic Acid (EGTA), MgCl_2_ and KCl were purchased from Fisher. Sterile colloidon (Nitrocellulose) 2% in amyl acetate and amyl acetate were purchased by Electron Microscopy Sciences Inc (Philadelphia, PA, US). d-Biotin was purchased from Avidity, LLC (Aurora, CO, US). Chloroform solutions of 1,2-Dioleoyl-sn-glycero-3-phosphocholine (DOPC) and 1,2-dioleoyl-sn-glycero-3-phosphoethanolamine-N-(cap biotinyl) (Biotinyl Cap PE), mini extruder, Nuclepore track-etch membrane (50 nm) from Whatman and filter support disks 10 mm from Whatman were purchased from Avanti Polar Lipids Inc (Alabaster, AL, US). High vacuum grease, Silicone elastomer base (184 Sylgrad) and the corresponding elastomer curing agent were manufactured by Dow Corning (Midland, MI, US). 2% Dimethyldichlorosilane in octamethylcyclooctasilane (PlusOne Repel-Silane ES) was purchased by GE Healthcare (Chicago, IL, US). Optical adhesive Norland 65 was purchased from Norland Products, Inc (Cranbury, NJ, US).

### Microbeads

Dry silica beads (dia. = 2.5 μm), streptavidin polystyrene beads (dia. = 0.51 μm, 1.0 % w/v; dia. = 0.82 μm, 1.0 % w/v; dia = 1.05 μm, 0.5 % w/v) and rabbit anti-6XHis polystyrene beads (2r = 0.61 μm, 0.1 % w/v) were purchased from Spherotech, Inc (Lake Forest, IL, US).

### Proteins

Unlabeled and labeled (biotin and TRITC) lyophilized tubulin from porcine brain were purchased from Cytoskeleton Inc (Denver, COL, US). Neutravidin was purchased from Life Sciences (St. Louis, MO, US). Casein powder, glucose oxidase from Aspergillus niger and bovine liver catalase were purchased from Sigma (St. Louis, MO, US). Mouse monoclonal anti-6xHis antibody was purchased from Abcam (Cambridge, MA, US). Biotinylated mouse anti-5xHis antibody was purchase from QIAGEN. Mouse anti Tubulin beta 3 antibody from BioRad (Hercules, CA, US). We used a 6xHis-GFP-labeled truncated human kinesin-1 heavy chain construct (amino acids 1-560; (26)) with an AviTag sequence susceptible to biotinylation added at the C-terminus (27). The protein was expressed in E. coli, purified using cobalt resin from Clontech Laboratories, Inc (now Takara Bio USA, Inc) (26). Biotin ligase BirA (Avidity LLC, Aurora, CO, US) was used to biotinylate Avitag-kinesin (28).

### Casein solution

Casein solution 20 mg/mL in 20 mM HEPES, 200 mM NaCl pH 8.8 was placed under gentle stirring either for 2-4 hours at room temperature or overnight at 4 °C. Solution was centrifuged at 110000 g, 4 °C for 20 min and supernatant was filtered through 150 mL rapid-flow filters, 50 mm in diameter and 0.2 μm pore size (Fisher). Concentrations were determined using the Bradford method, and solutions (8 to 13 mg/mL) were stored in 1 mL aliquots at −80 °C. When using GMPCPP microtubules, casein solution was prepared in KCl instead of NaCl because GMPCPP hydrolyzes faster in the presence of Na traces (29).

### Rectangular ridges

9 parts of silicone elastomer base were added to 1 part of the corresponding elastomer curing agent in a 50 mL screw-top cup conical tube (Corning, Tewksbury, MA, US) and were mixed gently by reversing the tube multiple times. The mixture was let undisturbed for 15 min. A chrome mask with the desired pattern of rectangular ridges (4 mm long, 2 µm wide, 1 µm tall and 10 µm apart) was placed face-up in a Pyrex borosilicate glass petri plate (Corning, Inc). The glass plate and the mask were placed next to a glass beaker containing 30 mL of 2% dimethyldichlorosilane solution in a vacuum chamber under a fume hood. A drop of 5 μL of 2% dimethyldichlorosilane (GE Healthcare) was added on the mask and hard vacuum was applied for 30 min. After releasing the vacuum, the elastomer solution was gently poured on top of the chrome mask in the glass plate and hard vacuum was reapplied for another 30 min to remove trapped air. To speed elastomer curing, the glass plate was placed at 105 °C for 1 hour. Using a razor blade, the cured elastomer on top of the chrome mask was removed and was used as a stamp to fabricate rectangular ridges on glass coverslips. Elastomer stamps were exposed to UV-plasma for no more than 4 sec and were then placed in a petri dish face-up. Dimethyldichlorosilane (5 μL of 2%) was placed on each stamp and let under hard vacuum for 30 min in the presence of a glass beaker containing 30 mL of 2% Dimethyldichlorosilane solution. Stamps were then placed gently face-down against ∼ 4 μl of optical adhesive (Norland 65) on glass coverlips and let for 5 min. To cure the optical adhesive, coverslips were exposed to UV light for 10 min. After gently removing the stamps, the cured optical adhesive on each coverslip was washed with 0.5 mL of methanol and dried in a fume hood. Coverslips with cured optical adhesive were used within 24 hours of preparation. A differential interference contrast microcopy image of the ridges is shown in Fig. S5A. Fluorescence microscopy image of microtubules (5% TRITC-tubulin) attached on the ridges are shown in Fig. S5B and Video S1.

### Supported Lipid Bilayers (SLBs)

Biotinylated supported lipid bilayers (SLBs) were prepared as in (30). In a round bottom flask 77 μL of 12.7 mM DOPC was mixed with 10 μL of 0.9 μM Biotinyl Cap PE. The flask was attached to a rotatory evaporator and the solution was thermally equilibrated in a water bath at 36 °C for 5 min. A hard vacuum was applied (30 min) to remove Chloroform, resulting in a lipid film. The lipid film was dissolved in 2 mL of buffer HNa100 (0.5 mM total lipid concentration) by rigorously vortexing the flask for 2 min at room temperature. The lipid suspension was subjected to at least four freeze-thaw cycles, followed by extrusion (11 passes) through 50 nm pores using a mini extruder (Avanti, Birmingham, AL, US) to form small unilamellar vesicles (SUVs). Chambers were assembled by pairs of detergent-washed and UV-plasma cleaned glass coverslips. SUVs were introduced in the chambers and incubated for 30 min. Chambers were placed in a humid box to prevent chambers from drying out. Chambers were washed by 3 × 100 μL with HNa100. Chambers were kept in a humid box until used within the same day.

### Kinesin attachment to polystyrene beads

Streptavidin-coated beads (10 μL) 0.82 μm in diameter were mixed with 89.5 μL of 2 mg/mL Casein in BRB80 pH 7.5 and bath sonicated for 1 min. Biotinylated anti-5xHis antibody (0.5 μL) was added and bead solution constantly rotated at 4 °C for at least 1 hr. d-Biotin was added to final concentration 2.6 μM and incubated for 1 hr at 4 °C under constant rotation. Bead solution was washed 3 times with 100 μL of 2 mg/mL casein by centrifugation at 6000 g, 4 °C for 5 min. Beads were stored at 4 °C under constant rotation and were used within a month. Beads (9 μL) were incubated biotinylated kinesin-1 (1 μL) under constant rotation at 4 °C for at least 4 hrs. To ensure single molecule interactions the kinesin-1 concentration was chosen so that no more than 1 out of 4 beads would interact with surface-immobilized non-biotinylated microtubules when assayed in the optical tweezers (31). For optical tweezers experiments, 2 μL of beads decorated with kinesin-1 were diluted to final volume of 50 μL with appropriate reagents (see final solution in Chamber preparation), and injected into the chamber. For surface-immobilized biotinylated microtubules, 8 μL of rabbit anti-6xHis polystyrene beads (0.61 μm in diameter) were mixed with 2 μl of casein 10 mg/mL and 2 μL of the appropriate concentration of kinesin-1. Beads were diluted 1:12 to final volume of 50 μl and injected into the chamber. For experiments comparing the effect of the bead’s size all three different sizes of beads where treated in parallel with the same stock of proteins and reagents and single molecule experiments were done the same day.

### Microtubule preparation

All tubulin solutions were 5 mg/mL in BRB80 pH 6.9 supplemented with 1 mM GTP. Unlabeled tubulin solution (20 μl) was mixed with 20 μL of biotinylated tubulin and 2 μL of TRITC tubulin. Tubulin solutions were centrifuged at 300000 g, 4 °C for 10 min using TLA-100 rotor (Beckman-Coulter). Tubulin in the supernatant was polymerized by incubating in the dark at 37 °C for 20 min. Taxol (1.75 μL of 2 mM) was then added, followed by 10 min incubation at 37 °C. Additional taxol (1 μL of 2 mM) was added and incubated for another 10 min at 37 °C. Polymerized tubulin solution was centrifuged at 40000 g, 25 °C for 20 min. Supernatant was discarded and the microtubule-containing pellet were resuspended by rigorous pipetting in 50 μl of BRB80 pH 6.9 supplemented with 40 μM taxol. Microtubules were kept at room temperature away from light and used within 5 days of preparation. For GMPCPP-stabilized microtubules, GTP and taxol were both excluded from the above protocol. Rather, 5 uL of 10 mM GMPPCP was added just before incubation of tubulin solution in a water bath at 37 °C, and tubulin polymerized for 30 min.

### Experimental chamber preparation

All solutions introduced in the experimental chambers and the necessary dilutions of stock solutions were prepared in buffer BRB80 pH 7.5. All glass coverslips were rectangular 22 × 40 mm and 1.5 mm thick (Fisher).

### Single-bead assay on a solid surface

Nitrocellulose-coated coverslips were assembled into flow chambers as described (32) and used within 24 hours of preparation. In experiments with rectangular ridges, one of the coverslips contains the imprinted optical adhesive. The volume of the experimental chambers was ≤ 20 μl. Solutions were introduced in the chamber in the following sequence: 20 μL of 0.05 mg/mL anti-tubulin antibody (BioRad) for 5 min; 50 μL of 2 mg/mL casein for 4 min; 4 × 25 μL of 125 nM microtubules supplemented with 2 mg/mL casein and 20 μM taxol for 4 × 1 min; washed with 100 μL of 2 mg/mL casein; 50 μL of final solution with kinesin-beads, 2 mM ATP, 2 mM MgCl_2_, 50 μM DTT, 20 μM taxol, 5 mg/mL glucose, 1500 units/mL of glucose oxidase and 0.2 units/mL of catalase. The open ends of the chamber were sealed with vacuum grease to prevent evaporation during the experiment.

### Single-bead assay on supported lipid bilayers (SLBs)

Solutions were introduced in chambers with SLB in the following sequence and for the following incubation times: 50 μl of 0.24 mg/mL neutravidin for 2 min; 4 × 25 μl of 1 μM 48% biotinylated microtubules supplemented with 20 μM taxol 4 × 1 min; 100 μl of 5 μM d-Biotin supplemented with 20 μM taxol; 50 μl of final solution with kinesin decorated beads (rabbit anti-6xHis polystyrene beads, dia = 0.61 μm), 2 mM ATP, 2 mM MgCl_2_, 50 μM DTT, 20 μM taxol, 5 mg/mL glucose, 1500 units/mL of glucose oxidase and 0.2 units/mL of catalase. The open ends of the chamber were sealed with vacuum grease.

### Three-bead optical tweezers assay

Nitrocellulose-coated flow chambers containing silica spherical pedestals (dia. = 2.5 μm) were prepared as described (32). Solutions were introduced in the chamber in the following sequence: 20 μL of 0.2 mg/mL anti-6xHis antibody (Abcam) for 5 min; 50 μL of 2 mg/mL casein for 4 min; 50 μl of kinesin-1 construct ≤1 nM supplemented with 2 mg/mL casein for 5 min; washed with 100 μL of 2 mg/mL casein; 50 μL of final solution containing 5 nM 48% biotinylated-microtubules, 2 mM ATP, 2 mM MgCl_2_, 50 μM DTT, 20 μM taxol (excluded when GMPCPP microtubules were used), 5 mg/mL glucose, 1500 units/mL of glucose oxidase and 0.2 units/mL of catalase. Before sealing the chamber with vacuum grease, 3 – 4 μL of streptavidin beads (dia. = 0.82 μm) diluted 1:30 in final solution without microtubules were introduced from one side of the chamber. The concentrations of kinesin used were such that no more than one out of four kinesin-decorated spherical immobilized pedestals interacted with microtubule dumbbells. The fraction of interacting spherical pedestals in the range of 0.2 to 1 as a function of kinesin concentration is much better described by a Poisson distribution that one or more (reduced χ^2^ = 0.012, ν = 3) rather than two or more (reduced χ^2^ = 7.2, ν = 3) kinesin dimers are interacting with a microtubule dumbbell (Fig. S1). The titration results compare well with single molecule titration for the single-bead assay (31). In many studies that use the single-bead assay the fraction of interacting beads for single molecule measurements ranges from 0.5 to 0.25 (4, 15, 16, 33). The concentrations of kinesin used in the current study were such that no more than one out of four kinesin-decorated spherical immobilized pedestals interacted with microtubule dumbbells which corresponds to fraction of interacting spherical pedestals ≤ 0.25.

### Optical Tweezers Experiments

Single-molecule interactions between microtubules and kinesin were recorded at 20 °C in a dual-beam optical-trap system using a 63 × water objective with numerical aperture of 1.2 (17), (34). For each trapped bead, the trap stiffness (pN/nm) and the system-calibration factor (pN/V) were determined by fitting a Lorentzian function to the corresponding power spectrum of their Brownian motion before microtubule attachment. Microtubule dumbbells were subjected to pre-tensile forces of 2 to 9 pN by moving the two laser beams apart. The trap stiffness of the individual laser beams for single-bead assays was 0.029–0.058 pN/nm and for three-bead assays 0.045 – 0.090 pN/nm. Higher stiffness than the single-bead assay was required in the three-bead assay to accommodate pretensile and kinesin-generated forces. A piezoelectric stage controller was used for position manipulation of single beads over surface-immobilized microtubules or microtubule dumbbells on top of spherical pedestals to scan for interactions. Data were filtered at 1 kHz, digitized with a sampling rate of 2 kHz and recorded using in-house software written in LabVIEW. Forces traces in Fig.1 were smoothed using Savitzky-Golay filter of 2^nd^ polynomial order and a 50-point half-window (Origin 2018b). To measure interactions between kinesin and microtubule dumbbells under zero pretensile force between the two beads (Fig. 2D, 2H, 2L), the laser that was trapping the minus-end streptavidin bead was shut off while kinesin was interacting with a dumbbell under pretensile forces. Only new processive kinesin runs, after the trap had been switched off, were quantified (Δ*t*, *F*_detach_) for this assay. After turning off the minus-end trap, thermal fluctuations of the microtubule when kinesin was not attached would often drive the microtubule away from the anchored kinesin. Therefore, the same procedure of detecting interactions under pretensile forces and then switching off the minus-end laser trap was repeated hundreds of times to collect a sufficient number (> 30) of interactions between kinesin and dumbbells in the absence of pretensile forces (minus-end trap off).

## Author Contributions

S.P., H. S. and E.M.O. designed the experiments. S.P. performed the experiments and analyzed the data. S.P., H.S. and E.M.O. wrote the manuscript.

## Acknowledgments

The authors would like to thank Aaron Snoberger for careful reading of the manuscript.

## Funding

NIH grant GM087253 to H.S and E.M.O, and NSF Science and Technology Center, CMMI: 15-48571to E.M.O.

## Notes

#### Summary of Updates

Single-bead assays with beads of different diameters to show that kinesin detaches faster for higher size beads due to the higher vertical component between the bead and the underlying surface.

