## Supporting Information for "Modulation of kinesin’s load-bearing capacity by force geometry and the microtubule track"

### Data Analysis

Using the system calibration factor (pN/V) experimental traces were converted to forces. In the three-bead assay the low velocity balance of forces on beads A and B and kinesin K is $\overrightarrow{F_A}(t) + \overrightarrow{F_B}(t) + \overrightarrow{F_K}(t) = 0$, or

$$\overrightarrow{F_K}(t) = -(\overrightarrow{F_A}(t) + \overrightarrow{F_B}(t)).$$

At times before kinesin interacts with the microtubule dumbbell, $\overrightarrow{F_K}(t < t_0) = 0$,

$$\overrightarrow{F_A}(t < t_0) + \overrightarrow{F_B}(t < t_0) = 0$$

and the forces of the two traps are equal and opposite to each other $\overrightarrow{F_A}(t < t_0) = -\overrightarrow{F_B}(t < t_0)$ and their magnitudes equal the pre attachment tensile force on the microtubule ($F_A = F_B = F_{pre\text{-}tensile}$). To reduce any offset or drift errors in the data collection the kinesin force is estimated to be

$$\overrightarrow{F_K}(t) = -[\overrightarrow{F_A}(t) + \overrightarrow{F_B}(t)] - [\overrightarrow{F_A}(t < t_0) + \overrightarrow{F_B}(t < t_0)]$$

$$\overrightarrow{F_K}(t) = -[(\overrightarrow{F_A}(t) - \overrightarrow{F_A}(t < t_0) + (\overrightarrow{F_B}(t) - \overrightarrow{F_B}(t < t_0)]$$

$$\overrightarrow{F_K}(t) = -[\Delta\overrightarrow{F_A}(t) + \Delta\overrightarrow{F_B}(t)]$$

Assuming that the bead-microtubule attachments and the microtubule itself are much stiffer than the two laser traps, the force generated by kinesin

$$\overrightarrow{F_K}(t) = -(k_A + k_B)\,\vec{d}(t) \qquad \text{Eq. S1 or}$$

$$\vec{d}(t) = -\overrightarrow{F_K}(t) / (k_A + k_B)$$

where $k_A$ and $k_B$ represent the stiffness values of the traps A and B and $\vec{d}(t)$ the displacement of the microtubule dumbbell by kinesin.

The magnitude of the average rate of change of the force $\Delta F_k(t)/\Delta t$ during kinesin's processive runs is:

$(F_k(t+\Delta t)- F_k(t))/\Delta t = (k_A + k_B) \cdot \Delta d(t)/\Delta t = (k_A + k_B) \cdot v_{av}$ Eq. S2

where $v_{av} = \Delta d(t)/\Delta t$ is the average velocity of kinesin for the time interval $\Delta t$. From Eq. S2 one can calculate the average velocity of kinesin.

From every dataset we calculate the average trace for kinesin's force production by averaging all kinesin's runs in the dataset. An example of the resulting average force trace is shown in Fig. S4A. The initial rising phase for $F < 3$ pN is approximated as a linear function of time and from the slope the loading rate dF/dt can be calculated. Then using Eq. S2 the average velocity $v_{av}$ is calculated. The distribution of the mean velocity and the corresponding box statistics, calculated as described above, for different pairs of single beads and surface immobilized microtubules and for different dumbbells are shown in Fig. S4B.

To every statistical quantity $q_i$, such as median-$\Delta$t and $v_{av}$, calculated for a dataset i, a statistical weight $w_i = n_i/n_{tot}$ is assigned, where $n_i$ represents the number of kinesin runs within dataset i and $n_{tot} = \Sigma n_i$ is the total number of kinesin runs from all datasets of the same assay. The weighted mean $\langle q \rangle_w$ and the weighted standard deviation $SD_w$ of the statistical quantity q are then calculated as follows:

$$\langle q \rangle_w = w_i q_i$$

$$S_w = \sqrt{\frac{\sum w_i (q_i - \bar{q})^2}{\frac{n_{t_i} - 1}{n_{t_i}} \sum w_i}}$$

**Stall forces:**

Stall forces were calculated by averaging within each dataset all force plateaus that lasted at least 0.1 s before detachment of kinesin (15, 35). To qualify as plateau, the standard deviation of the force should not exceed 5% of the average force value within the 0.1 s time window before detachment. In the case of the three-bead assay however a 10% threshold was used due to increased thermal fluctuations of the microtubule dumbbells.

**Statistical Comparisons:**

For statistical comparisons, the non-parametric Wilcoxon-Mann-Whitney test at the 0.05 confidence level was applied using either Origin Software 2018b or Studio R. When the sizes of the two compared datasets were different and the smaller size was characterized by smaller variance, the bigger size dataset was randomly truncated to the same size to avoid adverse effects on the comparison due to significant differences in the variances (36). When comparing more than two datasets the non-parametric Kruskal-Wallis test was used.

Video S1.

Movie of fluorescently labeled microtubules (5% TRITC-tubulin) attached on rectangular ridges shown in Fig. S1. Images were recorded at a rate of 1 fr/s for 21 seconds.

Video S2.

Cartoon animation showing the positional variability of a streptavidin-bead along the circumference of the microtubule in a dumbbell. The dumbbell is displayed in a cross-sectional view and the azimuthal angle $\varphi$ between the point of bead attachment and the protofilament that kinesin is interacting is indicated at every frame. The animation is not drawn to scale.

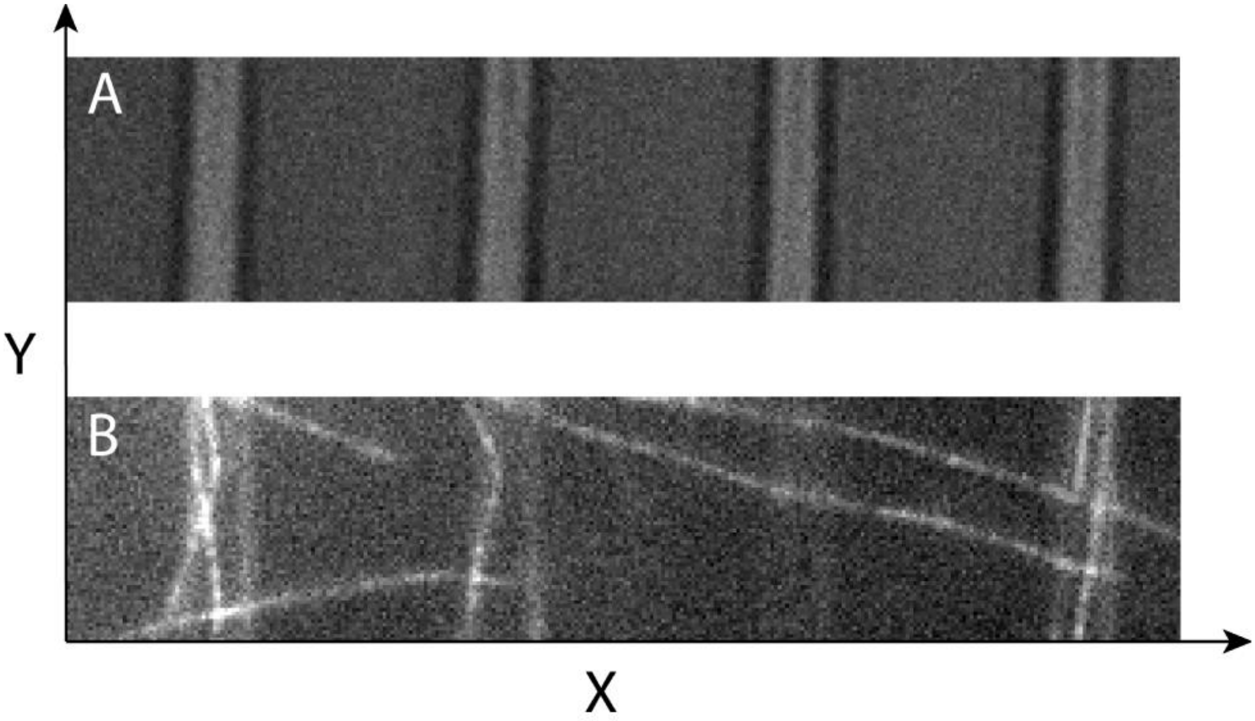


**Fig. S1. Microtubules suspended across rectangular ridges.**

(A) DIC (Differential interference contrast) microcopy image of rectangular parallel ridges (light gray stripes) with their long axis (4 mm) along the Y-direction and their short axis (2 μm) along the X-direction. Each ridge is 1 μm tall in the Z-direction and 10 μm apart from each other along the X-direction.

(B) Epifluorescence microscopy image of microtubules (5 % TRITC-tubulin) attached on the ridges shown in (A). The parts of the microtubules in-between the ridges due to lack of surface attachments are subjected to thermal fluctuations as can be seen in Video S1. Imaging was done on a Leica inverted microscope (DMI3000 B) using a 100 x oil objective from Leica and Metamorph imaging software. Further processing of the images was done using imageJ.

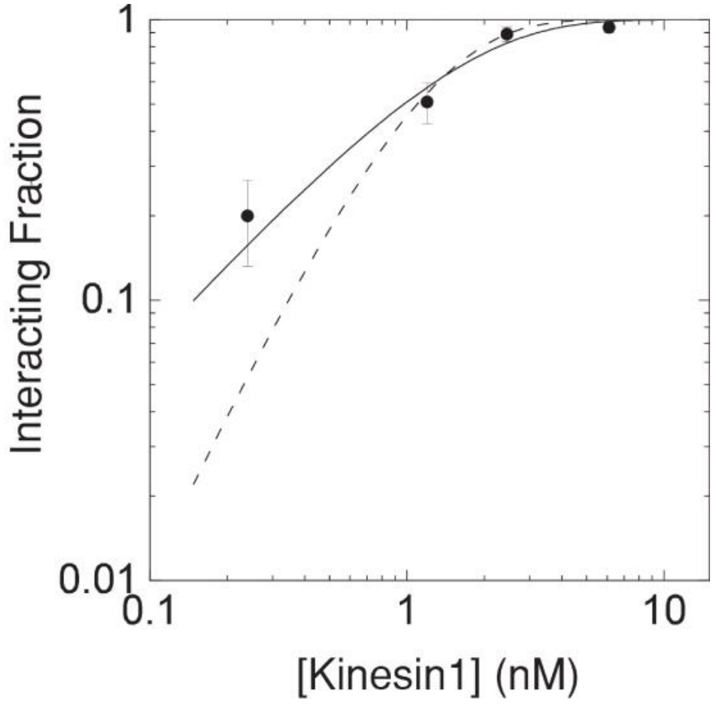


**Fig. S2. Single molecule titration for the three-bead assay.** The fraction of spherical pedestals that interact with a microtubule dumbbell in the range of 0.2 to 1 as a function of the kinesin concentration x used to decorate the spherical pedestals. For each kinesin concentration N=30 different pedestals were sampled in the same experimental chamber (scatter points). The solid and dashed lines represent fit of the data to the Poisson probabilities that at least one ($P(x) = 1 – \exp(-\lambda x)$) or at least two kinesins dimers ($P(x) = 1 – \exp(-\lambda x) - (\lambda x)\cdot\exp(-\lambda x)$) are interacting with the microtubule dumbbell, correspondingly. P stands for the fraction of interacting pedestals, $\lambda$ is a fitting parameter and error bars were calculated by the expression $[P(1-P)/N]^{1/2}$. The fit of the solid line ($\chi^2 = 0.012$, $\nu = 3$, $\lambda = 0.71$) is significantly superior compared to the dashed line (reduced $\chi^2 = 7.2$, $\nu = 3$, $\lambda = 1.5$).

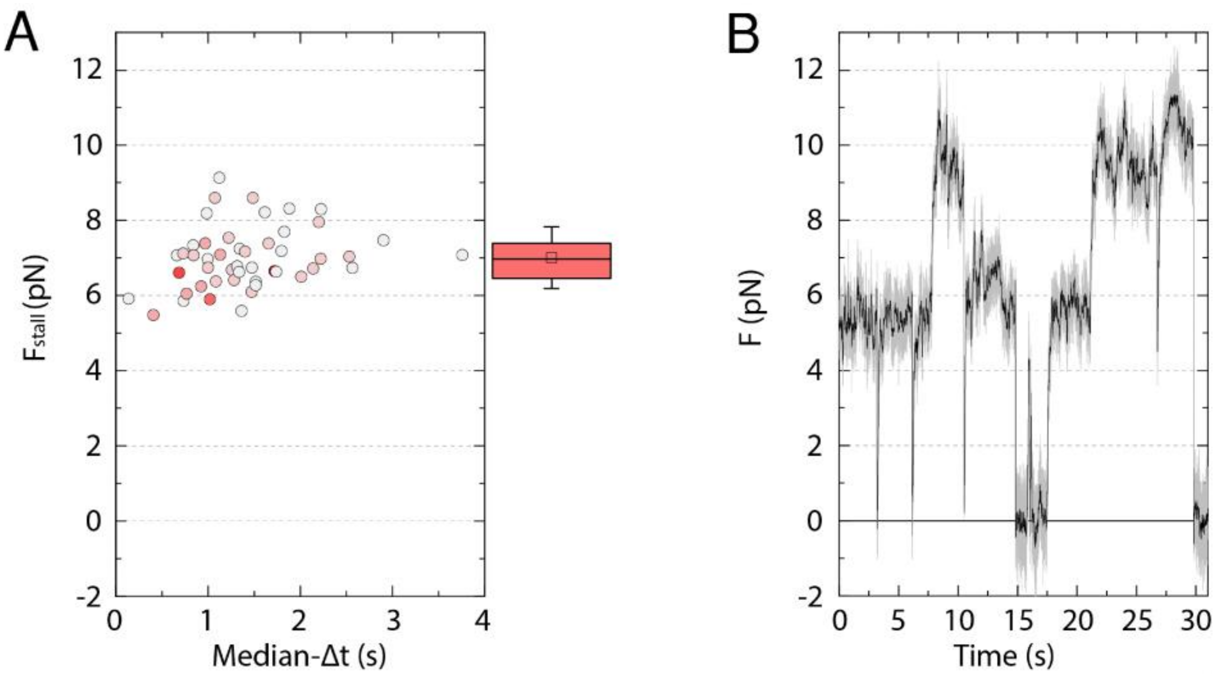


**Fig. S3. Stall forces for microtubule dumbbells do not scale with attachment durations.** (A) Scatter plot of the stall force $F_{stall}$ for each dumbbell as a function of the corresponding median-Δ*t*. In all cases, the fraction of interacting spherical pedestals was ≤ 0.25. Each scatter point corresponds to a different dumbbell and is the average of force plateaus that lasted for at least 0.1 ms before kinesin detachment. The shading of points scales with the corresponding number of force plateaus (statistical weight) for each point, with darker shading indicating a higher number. The value of $F_{stall}$ doesn't scale and correlates poorly with the corresponding median-Δ*t* ($r = 0.25$ and $p = 0.08 > 0.05$). The box statistics of $F_{stall}$ on the right reflects the symmetrical character of the distribution. (B) An example of a raw force trace (gray) and its smoothed version (black) when the fraction of interacting pedestals was 0.50. Unlike in Fig. 1D, not all the force plateaus are similar and the higher force plateaus ~ 10 pN are approximately twice as high as the lower ones ~ 5 pN due to the engagement of more than one kinesin molecules with the microtubule dumbbell.

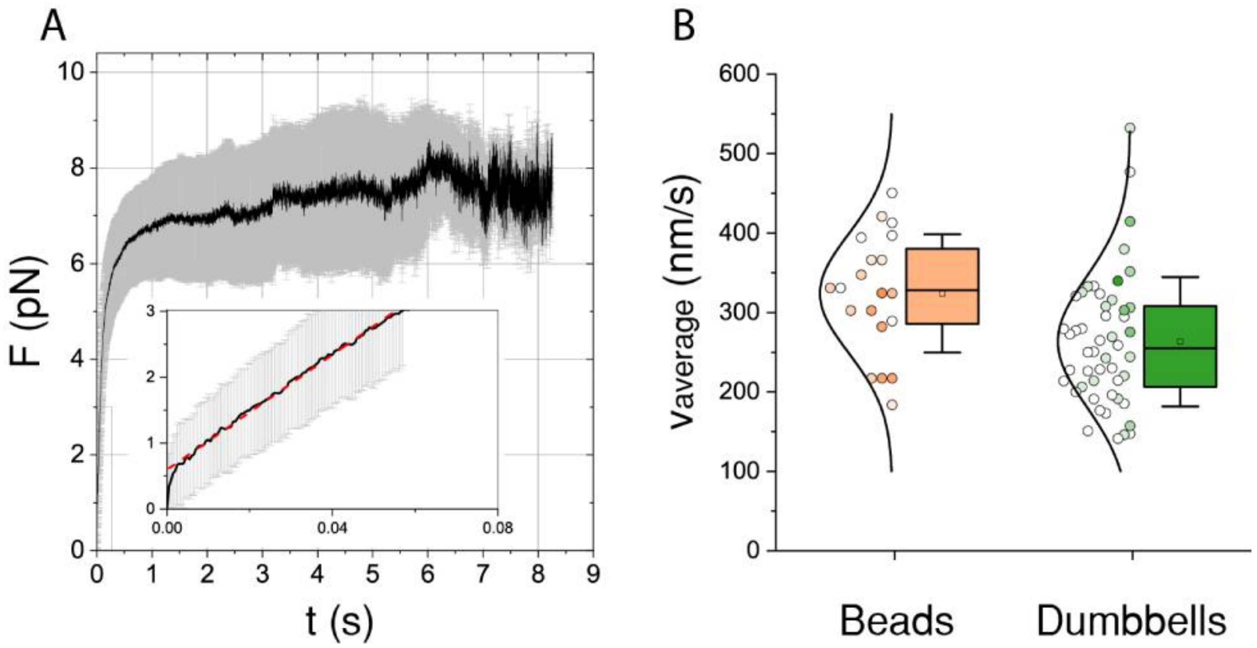


**Fig. S4. Average velocity of kinesin for the single-bead and three-bead assays.** (A) Examples of the average force trace and the corresponding standard deviation of all kinesin runs from a single dataset are shown by the black line and the gray error bars, respectively. A weighted linear fit (red dashed line) of the initial rising phase for F < 3 pN is shown in the zoom inset. Dividing the slope (pN/s) of the linear fit by the stiffness (pN/nm) the average velocity is calculated (Supporting Information Eq. S2. (B) The distribution of the average velocity and the corresponding box-statistics for 20 different pairs of single beads and surface immobilized microtubules (light brown color) and for 50 different microtubule dumbbells (green color). Each scatter point has been shaded based on its statistical weight, with darker shading indicating higher statistical weight (see Data Analysis).

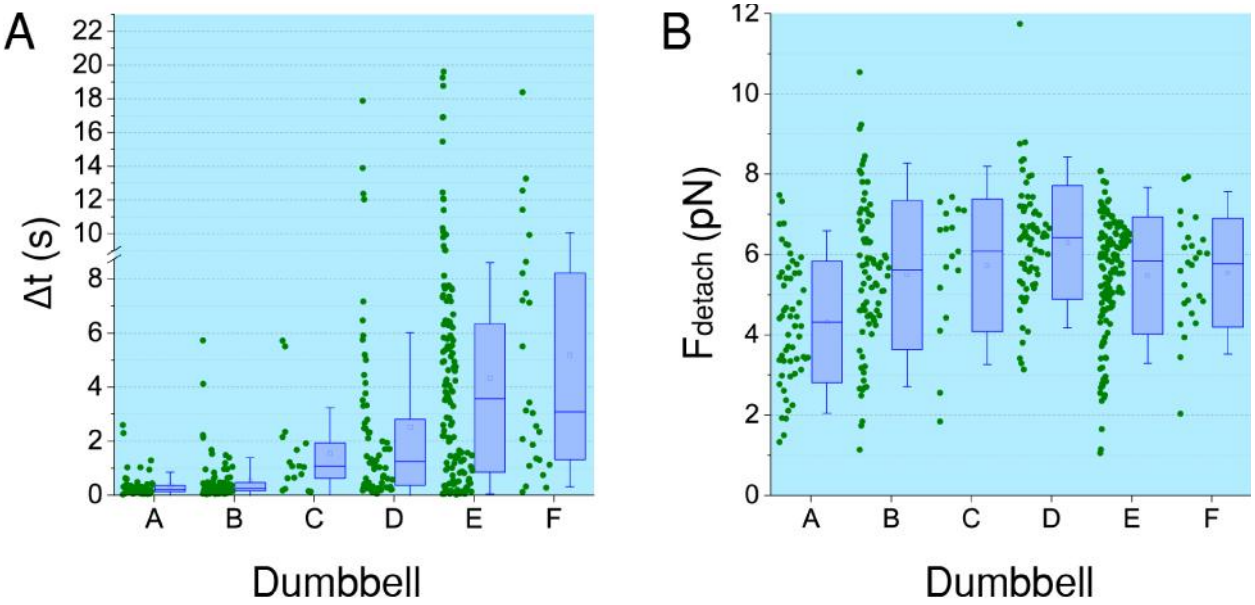


**Fig. S5. Distribution of attachment durations and detachment forces between kinesin and GMPCPP microtubules.**

Distribution and box statistics of (A) attachment durations Δt and (B) the corresponding detachment forces $F_{detach}$ for single molecule interactions between kinesin and GMPCPP microtubule dumbbells (“A” to “F”).

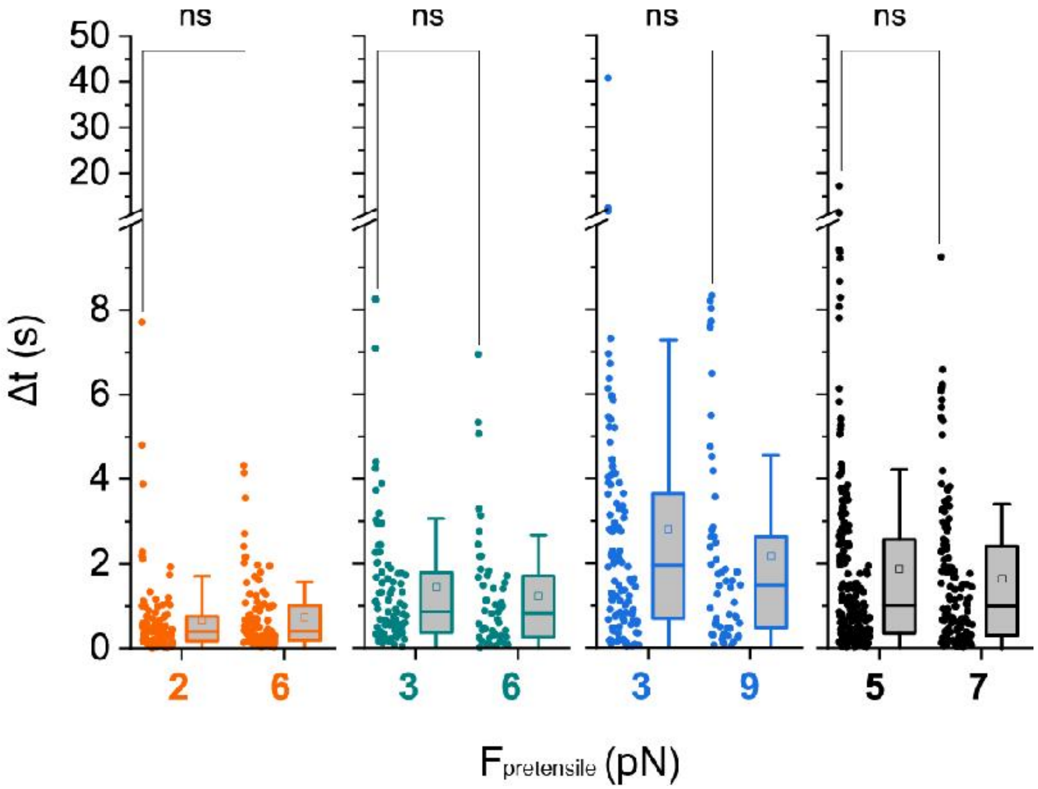


**Fig. S6. Attachment durations between kinesin and dumbbells under different pretensile forces.**

The distribution and box statistics of attachment durations Δ*t* between kinesin and 4 different microtubule dumbbells (different colors), each subjected to two different values of pre-tensile forces in the range between 2 and 9 pN. Statistical comparison showed that the magnitude of the pre-tensile forces for any given dumbbell doesn't seem to have an effect on the distribution of Δ*t* (Wilcoxon-Mann-Whitney test; $p>0.05$).

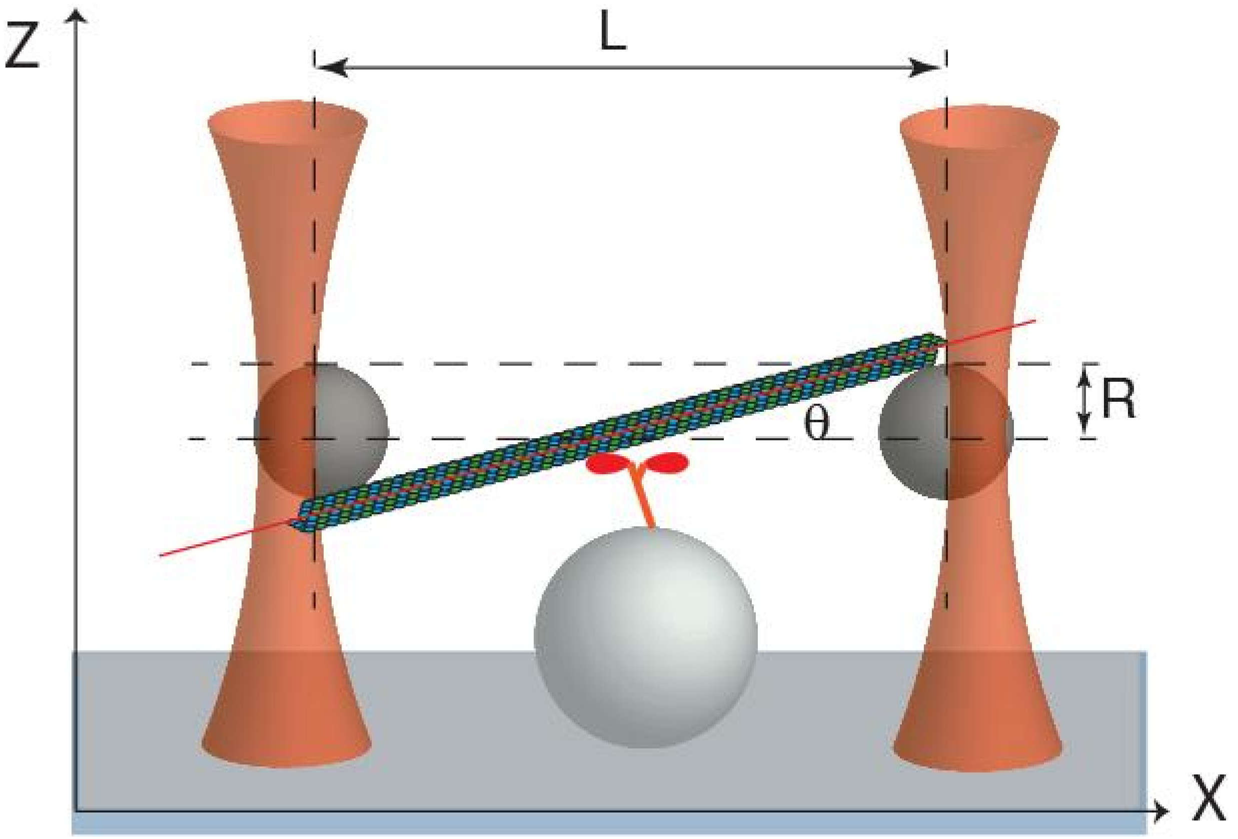


**Fig. S7. Z-force schematics for the three bead assay.** The 3D orientation of the microtubule within a dumbbell is quantified by the angle $\theta$ between the microtubule long axis (red line) and the horizontal x-axis. L (10 $\mu$m) is the distance between the two beads and R (0.41 $\mu$m) is the radius of each bead. The maximum displacement of a microtubule dumbbell in the z-direction as it is pulled by the surface immobilized kinesin happens for the configuration shown above, for which $\theta = \arctan(2R/L) = 4.7\ °$. Assuming that the microtubule is infinitely stiff, a force of 5 pN in the x-direction ($k_x \cong 0.05$ pN/nm) would correspond to a 2D displacement of $\Delta x \cong$ 100 nm and $\Delta z = \Delta x \cdot \tan\theta \cong 8$ nm. Since the stiffness of the laser trap in the z-direction is in general smaller than in the x-direction (37, 38), the force z-component of the force is not expected to exceed 0.4 pN.

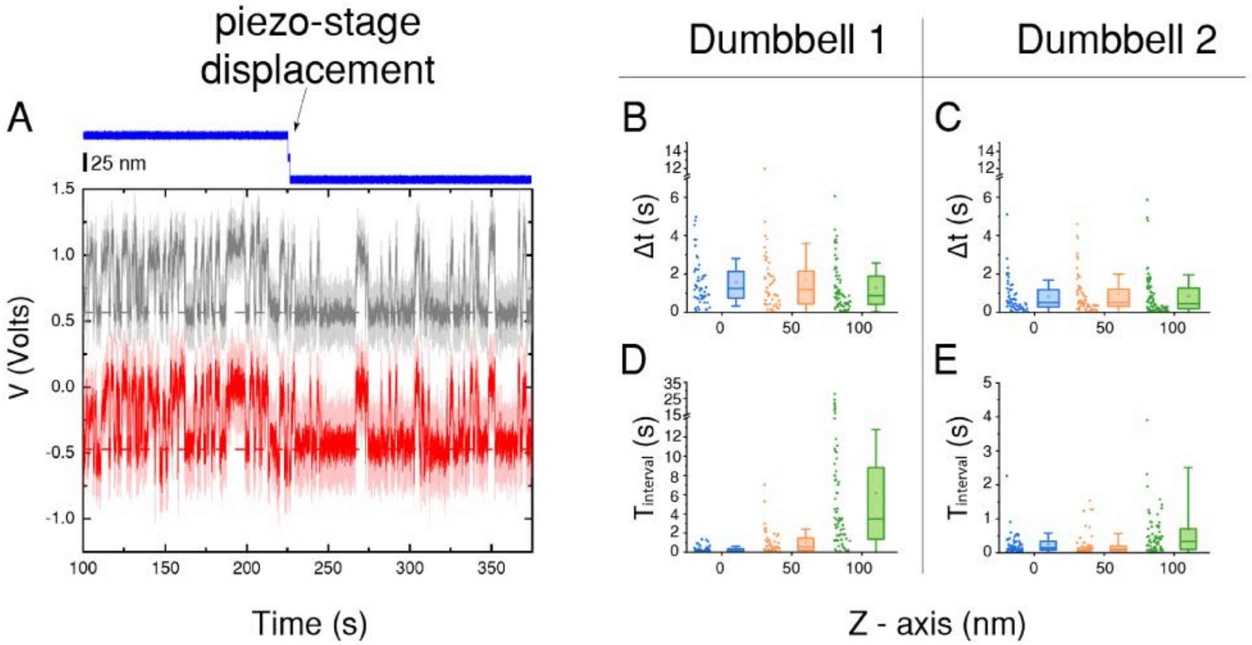


**Fig. S8. Variability in the z-position of microtubule dumbbells does not affect their attachment duration with kinesin**

(A) Sample of raw data traces (Voltage vs time) for the x-position of the two trapped beads (gray and red colors) during multiple interactions between a microtubule dumbbell and a kinesin molecule. The dark color traces are smoothed versions of the light color ones and the horizontal dashed lines indicate the positions of the two beads when the dumbbell is not attached to kinesin. The z-position of the piezo-stage is shown by the blue trace (top) and a total 50 nm displacement (scale bar 25 nm) of the dumbbell further away from the kinesin is indicated by the arrow. Note the decrease in the frequency of interactions after the z-displacement. (B) and (C) show plots of $\Delta t$ distribution for two different dumbbells at three different z-positions (three different colors) relative to the surface attached kinesin. The $\Delta t$ distributions for each dumbbell are not significantly different from each other at the 0.05 significance level (Kruskal Wallis test). (D) and (E) show the corresponding distribution of the time intervals $T_{interval}$ between successive kinesin-microtubule interactions in each case. The $T_{interval}$ distributions for each dumbbell are significantly different from each other at the 0.05 significance level (Kruskal Wallis test), in contrast to the corresponding $\Delta t$ distributions. Increasing the separation between the microtubule dumbbell and the surface attached kinesin affects only the frequency of their interaction and not the distribution of $\Delta t$.

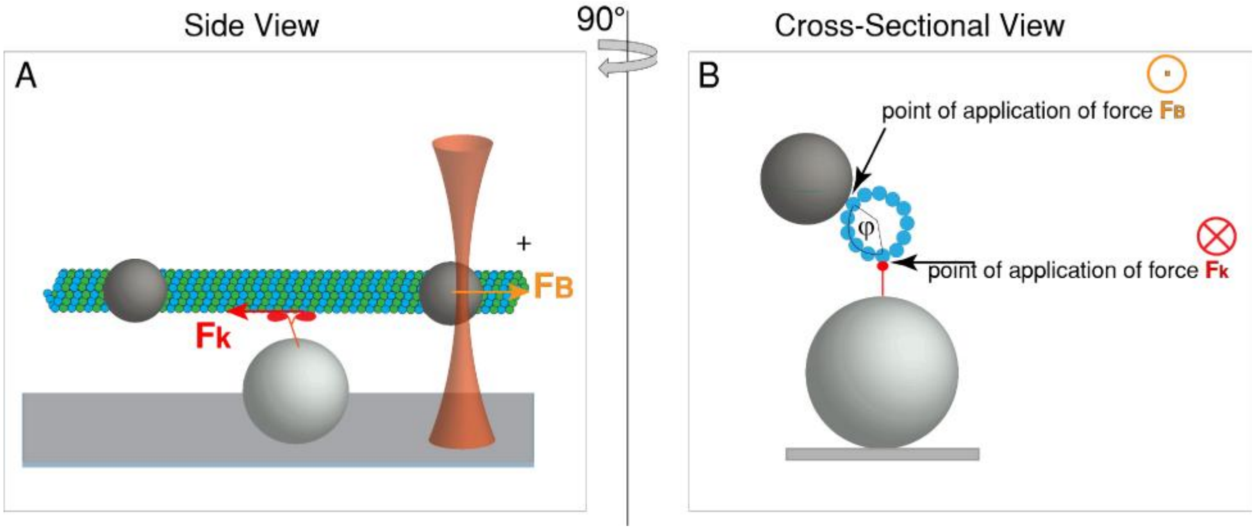


**Fig. S9. Cartoon representation (not drawn to scale) of the pair of opposing forces between kinesin and a microtubule dumbbell**. For simplicity we assume a three-bead assay in which only the plus-end bead is trapped by the laser. (A) The direction of kinesin's motion is towards the plus end (right) and therefore pulls the microtubule towards the opposite direction (left) by applying a force $F_k$ on the interacting protofilament. The stationary trap then develops an opposing force $F_B$ on the bead which is applied on the microtubule at the bead microtubule-attachment point. (B) Cross-sectional view of the same pair of forces as in (A). The relative azimuthal position $\varphi$ between the points of application of the opposing forces $F_k$ and $F_B$ is indicated. $F_k$ is directed vertically towards the back of the page ( $\otimes$ ) and $F_B$ toward the front ( $\odot$ ). For the single-bead assay the azimuthal separation of opposing forces is expected to be closer to 180$^\circ$ while it is more variable for the three-bead assay (Video S2).

**Table S1.**

**Different microtubule attachment strategies and substrates for single-bead assay (Fig. 2A)**

| | Median-Δt (s) | Representation in Fig 2A |
|---|---|---|
| Non-Biotinylated MTs immobilized via tubulin Ab on solid surface | 0.266 | a |
| | 0.425 | b |
| | 0.283 | c |
| | 0.261 | d |
| Biotinylated MTs immobilized via streptavidin on solid surface | 0.315 | e |
| Biotinylated MTs immobilized via streptavidin on Biotinylated lipid bilayer | 0.267 | f |
